# Translational insights into canine dorsal root ganglia cell types using cross-species comparisons

**DOI:** 10.64898/2026.04.14.718477

**Authors:** Leanne Jankelunas, Shamsuddin A. Bhuiyan, Hannah J. MacMillan, Thomas Cecere, Andrea Bertke, John H. Rossmeisl, William Renthal, Rell L. Parker

## Abstract

Chronic pain accounts for nearly half of owner-reported canine euthanasia decisions, yet dogs remain underutilized as a large-animal model for studying pain and developing translational therapeutics. Here, we present a canine dorsal root ganglion (DRG) cell atlas generated from six donors, representing five breeds, both sexes, and three spinal segments. Our dataset comprises 3,026 neurons and 11,734 non-neuronal cells and resolves 15 neuronal subtypes that map cleanly onto A- and C-fiber classes. We further identify eight major non-neuronal subtypes, including glial, vascular, and immune populations and characterize neuronal and non-neuronal expression of physiologically relevant neuropeptides, receptors, and ion channels. We identify region-specific differences in subtype composition between lumbar and sacral DRGs, with transcriptional programs suggestive of enhanced tactile-associated signaling in lumbar DRGs and heightened nociception-associated signaling in sacral DRGs. Cross-species comparisons reveal that canine DRG subtypes are broadly conserved with human and mouse, while also exhibiting canine-specific and canine–human shared molecular features relevant for translation. Together, this atlas serves as a valuable resource for understanding canine sensory neurobiology, comparing DRG organization across mammals, and leveraging dogs as a translational model for pain research and therapeutic development.

## Introduction

Pet dogs often suffer from chronic pain and itch disorders, for which limited therapeutic options are available [17; 38]. Estimates of pain prevalence in dogs include 20% of dogs treated as outpatients and 22% of dogs in an intensive care unit setting. Dog owners report pain and/or suffering as the main cause for euthanasia 49% of the time [39; 40; 42]. Understanding the mechanisms underlying pain and the primary sensory neurons contributing to nociception is therefore imperative for improving dog health.

While under-utilized in pain research, dogs have a long history of serving as models for human disease due to their shared environment, diet, and longer lifespan when compared to other translational models [41]. Humans and dogs share many aspects of anatomy and pathophysiology; therefore, dogs may be an excellent translational model for pain [24; 28; 29]. A deeper understanding of the canine primary sensory neurons in the dorsal root ganglion (DRG) may allow for improved insight of DRG physiology across species. To better determine the potential for dogs as translational research models of pain, the generation of a canine DRG atlas is needed.

The DRG receives sensory information related to the internal and external environment that is transmitted to the spinal cord and brain for conscious and unconscious perception. DRG neurons are specialized for mechanosensation, proprioception, thermosensation, visceral sensation, pruritoception, and nociception [10]. The two main functional neuronal groups, A fibers and C fibers, are classified by myelination, axonal size, and conduction velocity [1; 43]. DRG neurons can be further subclassified based on gene expression and electrophysiologic properties [27; 45]. The DRG also contains supporting cells such as satellite glia and resident immune cells, which maintain the neuronal microenvironment [21; 32].

To date, the primary sensory neurons within the DRG have been characterized in mice, rats, guinea pigs, pigs, macaques, and humans and key interspecies differences and similarities have been described [7; 25]. For murine models, often utilized in pain studies, the volume of cells within the DRG and size of the DRG is significantly less than human counterparts [20; 45; 56]. Interspecies differences also include shifts in cell-type–specific expression of key somatosensory molecules, including neuropeptides such as *Tac3* and ion channels such as *Trpm8* [7; 22; 25; 37; 44]. This illustrates the need for additional translational models of nociception.

The goal of this study was to determine the cell types in the canine DRG, including both neurons and non-neurons, and to compare the canine DRG to the available human, murine, primate, and swine DRG atlases. Toward this end, we developed a protocol to collect canine DRGs and perform 10x Genomics snRNA-seq. We conducted cross-species and cross-spinal segment comparisons, identifying conserved and divergent cell types as well as differences in key therapeutic targets. This canine DRG cell atlas can be used to better understand the physiologic properties of touch and nociception across species and potentially serve as a resource for the development of new targets for therapeutic interventions for dogs.

## Methods

### DRG collection

Client-owned dogs between the ages of 1 year to 10 years who were being euthanized for medical reasons were considered for inclusion. Owner consent for collection of research samples was obtained in all cases. Potential donors were screened based on a review of available medical records for historical neurologic conditions, painful conditions affecting the pelvic limbs, or signs on physical examination that could be associated with underlying neurologic or orthopedic diseases. Dogs were excluded if there was a reported history of pelvic limb lameness or if there were any abnormalities on examination or a history of other potentially painful neurologic or orthopedic conditions of the lumbosacral region. When a full examination was not available, the potential donor’s history was reviewed.

Samples were collected within two hours of euthanasia based on AVMA-approved guidelines at the discretion of the overseeing clinician. Donors ultimately selected for inclusion of DRG tissue processing and atlas generation had to have no evidence of obvious disc degeneration or compressive lesions in the spinal cord and intervertebral foramina on necropsy. For the 10x library, we ultimately included three dogs consisting of 2 females and 1 male (**Table S1**).

### Sample Collection and Tissue Processing

Lumbar and sacral DRG were collected. DRGs were trimmed after collection using a dissecting microscope to remove excess tissue from the DRG. The individual DRGs were then snap-frozen in liquid nitrogen and individually stored in collection tubes at −80°C until samples could be further processed.

### DRG nuclei isolation

Nuclei isolation was performed as indicated in the 10x Genomics (Pleasanton, CA) CG000505 Chromium Nuclei Isolation Kit Protocol (1000494). Each DRG was sectioned on glass Petri dishes over dry ice and chopped into pieces no more than 0.25 mm using a razor blade. The DRG cells were then dissociated using the lysis buffer and pestle pulverizer provided in the 10x Genomics Chromium Nuclei Isolation kit and homogenization was performed until the entire sample could pass through a 1000 μL pipette tip without clogging. The sample was then moved to the lysis buffer for additional processing to obtain single nuclei. Once lysed, each sample had debris removed and was washed using the provided reagents in the 10x Genomics Chromium Nuclei Isolation kit and a cooled swinging bucket centrifuge. To reduce the degree of RNA degradation, each individual DRG was fully processed to isolate the nuclei before processing another sample. Before sequencing, the samples of isolated nuclei were evaluated using Trypan blue stain and manually counted with a hemocytometer to determine the concentrations of nuclei in each sample before loading into a 10x Genomics Chromium controller.

### Preparation of single-nuclei RNA-sequencing libraries

After nuclei isolation, single nuclei processing was performed using the Genomics 10x CG000731 Single Cell 3’ Reagent Kits v4 (1000269) as indicated in the manufacturer’s users guide and illustrated in **Figure 1**. A master mix containing the nuclei of each individual DRG was generated with an aim for sequencing 4,000 to 10,000 nuclei per DRG sample. This variation depended on the estimated nuclei yield from the nuclei isolation step, which was determined based on an automated cell counter. This master mix was then utilized for GEM generation and barcoding via a Chromium^TM^ Controller (10x, Pleasanton, CA). The samples underwent post-GEM-RT Cleanup, cDNA Amplification (11–12 cycles with v4 based on initial nuclei concentrations), and gene expression library construction as detailed in the manufacturer’s users guide. The gene expression libraries were generated by annealing one Dual Index Plate TT Set A label to each sample, which allowed for individual identification of each sample during sequencing. Sample index PCR was performed with 14 cycles.

**Figure 1:**
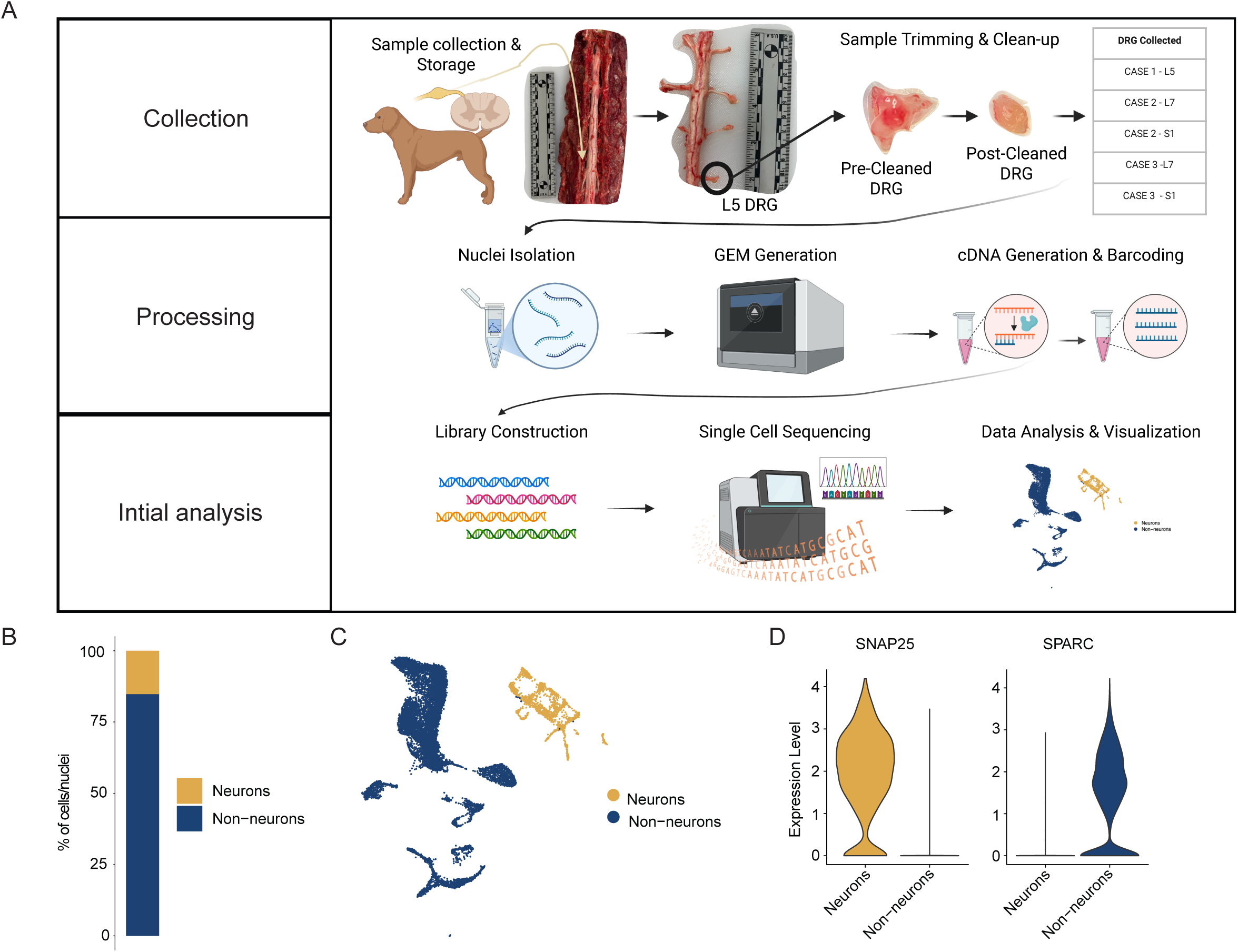
Methodology and preliminary classification into neuronal and non-neuronal cell populations. A) A stepwise illustration of the DRG collection, processing, library generation, and initial data analysis. During collection, a dorsal laminectomy was performed on each donor dog to visualize the spinal cord, nerve roots, and DRG. The DRG were isolated from as many levels in the lumbosacral region as possible. The individual DRG were trimmed to generate the post-cleaning DRG, which were then frozen. During processing, nuclei isolation was performed; then, the libraries were generated for initial analysis. B) Stacked bar plot displays the proportion of neuronal and non-neuronal cells/nuclei (color) in the DRG canine atlas. C) UMAP displays the non-neuronal and neuronal cells/nuclei (14,760 cells/nuclei total). UMAP is colored by neuronal or non-neuronal annotation. D) Violin plot displays log-normalized gene expression of the neuronal marker, *SNAP25*, and the non-neuronal marker, *SPARC.* Gene expression distribution is split by neuronal and non-neuronal annotations.

At two time points, after cDNA amplification and after post library construction, quantification of generated libraries for quality control was performed by Qubit (Picogreen) Assay Kit at the Fralin Life Sciences Institute Genomics Sequencing Facility. Profiling of the generated libraries was performed by High Sensitivity D5000 ScreenTape (Agilent TapeStation Software 4.1.1, Santa Clara, CA) and sequencing was performed by Illumina NovaSeq SP v1.5 200 cycle for (libraries 1 & 2) and Illumina NextSeq 2000 v3 P3 100 cycle (libraries 3–7) (Illumina, San Diego, CA) following the manufacturer’s guidelines at the Fralin Life Sciences Institute Genomics Sequencing Facility.

### Single nuclei RNA-seq processing and integration with Flash-seq

The construction of our canine atlas followed a similar protocol as Bhuiyan et al., 2024 and Bhuiyan et al., 2025 [6; 7]. Raw FASTQs from all 10x snRNA-seq datasets were aligned to the canine reference genome to generate counts matrices, followed by ambient RNA correction and empty droplet removal using CellBender (“remove-background” function). We then combined all 10x snRNA-seq datasets into a single Seurat object, split the merged object by dataset (or sequencing center), and performed normalization and variable feature selection using the vst method, while excluding mitochondrial genes (prefix “MT-”). We identified integration features and anchors across subsets and integrated datasets using Seurat CCA to generate an “integrated” assay. The integrated object was scaled and analyzed by PCA; PCs were selected using the same 90% cumulative variance and 5% marginal variance criteria, followed by computation of UMAP and t-SNE embeddings, construction of a shared nearest-neighbor graph, and clustering at resolution of 1.5. For additional information on the Flash-seq data set, which included nine dogs, (four males and five females) see (Fernandez et al.; under review)[15]. For quality control, we generated study-stratified violin plots of nFeature_RNA, nCount_RNA, and percent.mt and visualized marker expression on UMAP and t-SNE using broad cell-type markers (*SNAP25, RBFOX3, SPARC, MPZ, RBFOX1, PTPRC*); clusters consistent with doublets were removed when they showed concurrent enrichment for both neuronal and non-neuronal marker programs.

We then subclustered neurons and non-neurons separately, following the same clustering parameters as described above, with the key exception that the neuronal subclustering included integration with the Flash-seq data. Subtypes were annotated using the marker genes listed in **Figures 2B** and **3B**.

**Figure 2:**
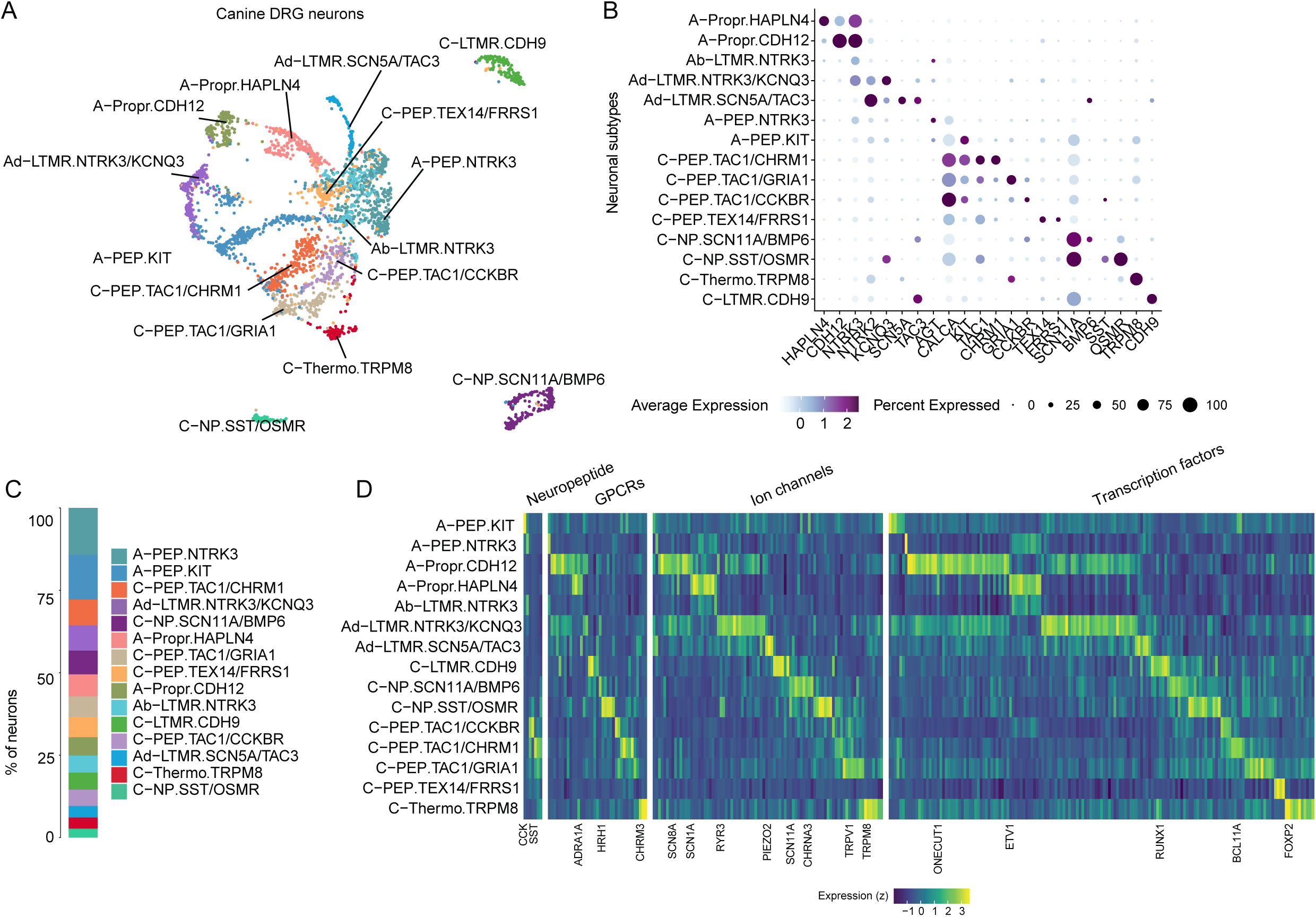
Transcriptional diversity of 15 canine DRG neuronal subtypes. A) Canine DRG neuronal atlas (3,026 cell/nuclei) projected into UMAP space. Cells/nuclei are colored by their final cell type annotations. B) Dot plot displays subtype-specific marker gene expression in DRG neuronal subtypes. Dot size indicates the fraction of cells/nuclei expressing each gene, and color indicates average log-normalized scaled expression of each gene. C) Stacked bar plot displays the proportion of each neuronal subtype (color) in the canine DRG neuronal atlas. D) Heatmap displays genes classified as neuropeptides, GPCRs, ion channels, and transcription factors in each neuronal subtype relative to all other neuronal subtypes (log2FC > 0.5, adj. P-value < 0.05). For visualization, only the top 15 genes (or less) are displayed per subtype.

**Figure 3:**
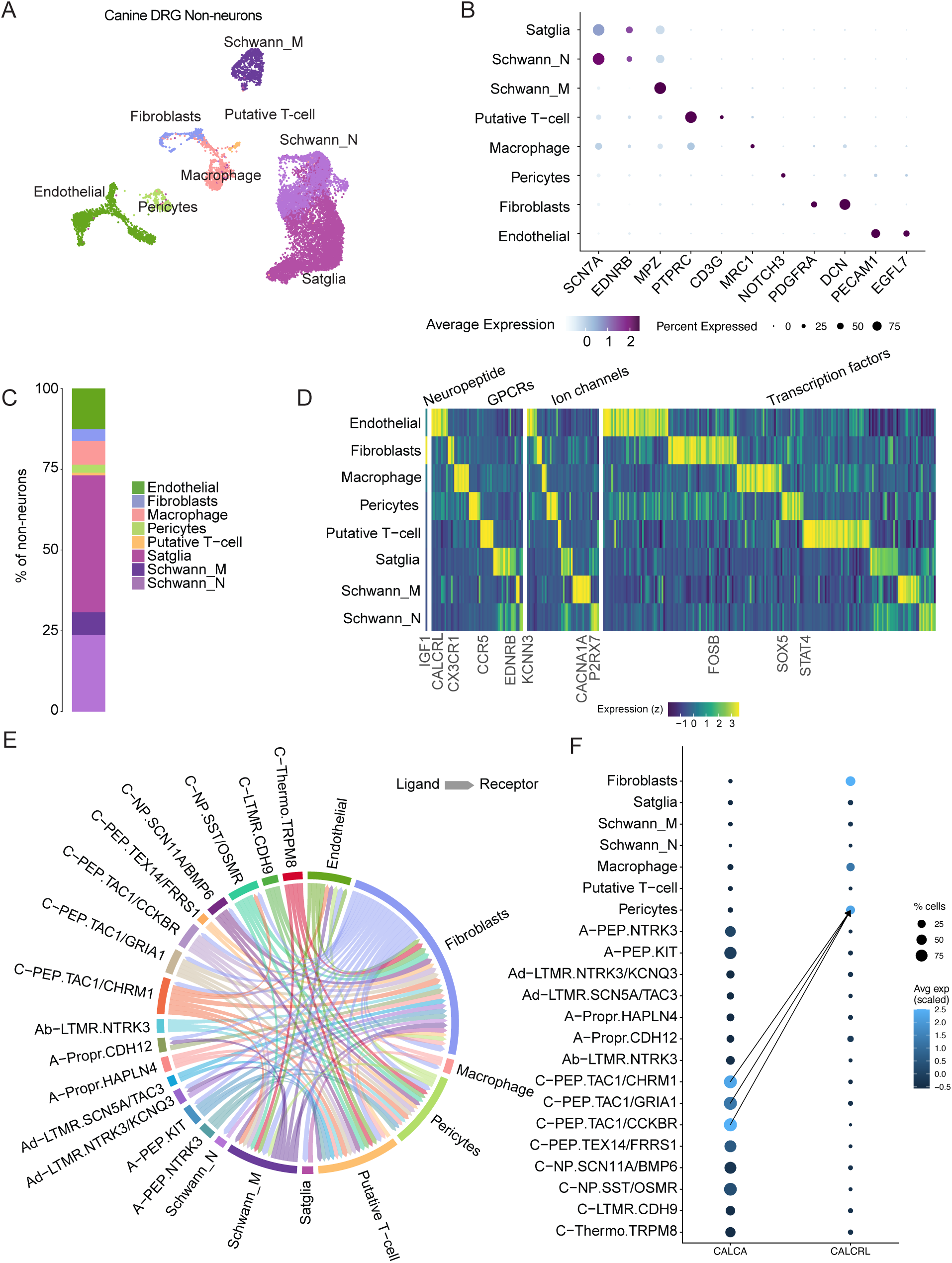
Transcriptional diversity of eight canine DRG neuronal subtypes. A) Canine DRG non-neuronal atlas (11,734 nuclei) projected into UMAP space. Nuclei are colored by their final cell type annotations. B) Dot plot displays cell-type-specific marker gene expression in DRG non-neuronal subtypes. Dot size indicates the fraction of nuclei expressing each gene, and color indicates average log-normalized scaled expression of each gene. C) Stacked bar plot displays the proportion of each non-neuronal subtype (color) in the canine DRG neuronal atlas. D) Heatmap displays genes classified as neuropeptides, GPCRs, ion channels, and transcription factors in each non-neuronal subtype relative to all other non-neuronal subtypes (log2FC > 0.5, adj. P-value < 0.05). For visualization, only the top 15 genes (or less) are displayed per subtype. E) Chord diagram displays ligand receptor predicted interactions between neuronal and non-neuronal subtypes. Only subtypes with an aggregated rank < 0.01 are displayed (see Methods). F) Dot plot displays the gene expression of ligand *CALCA* and its receptor *CALCRL* in individual canine DRG neuronal and non-neuronal subtypes. Dot size indicates fraction of cells/nuclei expressing each gene, and color indicates average log-normalized scaled expression of each gene. The arrows link the top 3 (ranked by aggregated rank) subtype pairs for each ligand–receptor interaction.

### Ligand-receptor analysis

LR pair analysis was performed using LR analysis framework (LIANA). The canine DRG non-neuronal and neuronal atlases were combined. We then ran LIANA with default settings using their human consensus database, and we filtered LR interactions with an aggregated rank of <0.01. Both the ligand and receptor genes needed to be expressed in at least 5% of cells/nuclei across the cell types used for the analysis. For each ligand–receptor interaction, the top three cell-type pairs ranked by the aggregated rank generated by LIANA were used for visualization.

### Cross-species comparisons

To enable cross-species comparisons, canine genes were mapped to their human orthologs using Ensembl BioMart (v109), following a strategy similar to Bhuiyan et al., 2024. Briefly, genes were first converted to 1:1 human orthologs where available. For genes with 1 to many or many to many relationships, we retained the ortholog with the highest orthology confidence score; where multiple candidates remained or where confidence scores were unavailable, the ortholog with the highest percentage identity was selected. Canine genes without a known human ortholog were retained in the counts matrix but excluded from downstream integration steps. Following ortholog mapping, we performed label transfer by anchoring the canine nuclei to a reference human DRG atlas using Seurat v4. Transfer anchors were identified using canonical correlation analysis (FindTransferAnchors function with the reduction argument set to “cca”), and cell type labels were transferred using the TransferData function. Variable features were identified from the combined dataset, followed by PCA and UMAP for visualization. Cells with low prediction confidence (score < 0.5) were excluded from downstream analyses.

## Results

### A protocol to collect canine DRGs for 10x Genomics single nuclei RNAseq

DRG samples were initially collected from three donors (**Table S1**) and immediately flash frozen, which led to an excessive amount of debris in the sample. Therefore, the protocol was altered to add a step to remove excess tissue from the DRG prior to freezing (**Figure 1A**). DRG nuclei isolation was challenging, and when the initial samples (A1 and B1) were processed, they were placed immediately in the 10x Genomics lysis reagent with pestle. Dissociation time was excessive, taking approximately 45 minutes per sample, leading to poor nuclei yield. Subsequent DRG were chopped into 1–2 mm pieces before being added to the lysis reagent, which substantially improved yield. For later samples, each DRG went through the nuclei isolation step to a pre-defined stopping point before processing the next DRG. To further increase nuclei capture, the last set of samples was processed using a swinging bucket centrifuge. These sequential improvements increased nuclei recovery, though the level of DRG (and therefore relative size) was also an important factor for total nuclei yield. All samples were then prepared uniformly for sequencing following the 10x CG000731 Single Cell 3’ Reagent Kits v4 protocol.

### The canine DRG atlas identifies 15 neuronal subtypes

Once the snRNA-seq data were generated, reads were aligned to the canine reference genome and counts matrices were produced. In parallel, scRNA-seq data were independently generated using Flash-seq (Fernandez et al.; under review)[15]. We constructed a canine DRG cell atlas by integrating the 10x Genomics snRNA-seq and Flash-seq datasets, leveraging the large number of nuclei captured by snRNA-seq and the greater transcript coverage, including cytoplasmic RNA, provided by Flash-seq. The cohort of six donors included after quality control represents five breeds, both sexes and three spinal levels. For the 10x snRNA-seq libraries, one L5, three L7, and two S1 DRG were sequenced. These were obtained from three different dogs. Altogether, this resulted in a canine DRG atlas of 3,026 neurons and 11,734 non-neurons (**Figure 1B–D**).

Across the 3,026 neurons, we observed 15 neuronal subtypes (**Figure 2A**). We assigned each subtype as A- or C-fiber using *NEFH and NEFM*, which are predefined markers set for fast-conducting, myelinated neurons (**Figure 2B**) [5; 7; 26]. Seven subtypes were annotated as A-fibers and eight as C-fibers. We then annotated neuronal subtypes using a standardized scheme from the human reference atlas generated by the NIH PAIN PRECISION Network: primary class (A or C), functional family based on marker genes from previous studies, and one or two discriminative marker genes (**Figure 2B**) [18; 47].

The seven *NEFH*-high A-fiber subtypes include two *PVALB+* proprioceptors (A-Propr), one *NTRK3-*high/*NTRK2*-low Ab-low threshold mechanoreceptor (Ab-LTMRs), two *NTRK3-*low/*NTRK2*-low Ad-LTMRs, and two *CALCA+* peptidergic nociceptor subtypes (A-PEP) [2; 11; 45; 49]. The eight *NEFH*-low C-fiber subtypes include a *TRPM8+*/*FOXP2*+ thermosensor (C-Thermo.TRPM8) [4], a *CDH9+*/*TAFA4+* LTMR (C-LTMR.CDH9), four *CALCA+/TAC1+* peptidergic nociceptor (C-PEP), and a *SCN11A+* non-canonical peptidergic nociceptor (C-NP). All 15 neuronal subtypes were detected in both the 10x Genomics snRNA-seq and Flash-seq datasets (**Figure S1A–C**). A-fiber neurons, however, were frequently underrepresented in the Flash-seq data, likely reflecting challenges related to their large cell size during cell sorting (**Figure S1B**). Interestingly, we observed that A-PEPs were the most abundant population in the canine DRG (**Figure 2C**), which differs from human DRGs, where C-PEPs were the most abundant [6].

Integration of Flash-seq, which provides greater transcript coverage, with snRNA-seq, which captures a larger number of nuclei, enabled the identification of genes enriched in each neuronal subtype relative to all other neurons. We found that the 15 subtypes were enriched (log_2_fc > 0.5, adjusted [adj.] p-value < 0.05 relative to other neurons) for 975 genes (**Table S2; Figure 2D**). These included therapeutically relevant gene families such as 7 neuropeptide (e.g., *CCK*), 37 GPCRs (e.g., *HRH1*), 86 ion channels (e.g., *CACNA1E*), and 159 transcription factors (e.g., *FOXP2*).

Collectively, these data describe cell-type-specific expression genes expressed at low levels, achieved through approximately 5 log-fold greater transcriptomic depth than the individual 10x snRNA-seq and Flash-seq datasets.

### Eight non-neuronal subtypes were identified in the canine DRG

As with human and mouse DRGs, non-neuronal canine DRG cells play crucial roles in regulating the neuronal microenvironment, modulating neuronal physiology through direct interactions and responding to changes in the external environment. In the canine DRG atlas, we observed eight major non-neuronal cell types (**Figure 3A**). These cell types include *PECAM1+* and *EGFL7+* endothelial cells, *DCN+* and *PDGFRA+* fibroblast cells, *EDNRB+* satellite glia cells, *MPZ+/MPB+* myelinating Schwann cells, *SCN7A+* non-myelination Schwann cells, and *PTPRC+/MRC1+* macrophages and *PTPRC+/CD3G+* T-cells (**Figure 3B**). While only the 10x snRNA-seq data set contained non-neuronal cells, we observed that all non-neuronal cell types were observed across all libraries with exception of the rarest population (T-cells; **Figure S2A**). The percentage of cell types across libraries is shown (**Figure 3C**).

We next examined genes relevant to non-neuronal physiology, including those encoding neuropeptides, receptors, and ion channels. We found that the eight subtypes were enriched for 3,204 genes (log_2_fc > 0.5, adjusted [adj.] p-value < 0.05 relative to other neurons) (**Table S3; Figure 3D**). These included therapeutically relevant gene families such as a single peptide (e.g., *IGF1*), 56 GPCRs (e.g., *CALCRL*), 44 ion channels (e.g., *P2RX7*), and 204 transcription factors (e.g., *SOX5*).

### Ligand-receptor analysis of the canine DRG cells

After observing the expression of ligand and receptor genes in both neuronal and non-neuronal subpopulations, we performed ligand-receptor (LR) interaction analysis between the canine neuronal and non-neuronal atlases to characterize putative interactions between subtypes. These analyses revealed over 200,000 LR interactions (**Figure 3E**), with fibroblasts, putative T-cells, and pericytes having the most predicted interactions. The large number of predicted fibroblast interactions remained even after controlling for the number of cells/nuclei per subtype (**Figure S2B**). Among LR analysis results, interactions with well-established roles in inflammation, including predicted interactions between *CALCA* (expressed across all three C-PEP subtypes) and *CALCRL* in pericytes, were observed (**Figure 3F**).

Together, our canine DRG atlas provides a preliminary cellular and molecular map of the canine DRG, resolving 15 neuronal and eight non-neuronal subtypes. The atlas uncovers previously unrecognized A-fiber cell types, rare C-fiber populations, and novel neuro-immune and neuro-glial signaling pathways that lay a foundation for studying canine somatosensory biology and pain mechanisms. We have made these atlases publicly available as an additional Pain-seq web resource: https://painseq.shinyapps.io/canine_drgs/.

### DRG subtype differences between canine spinal segments

Different body regions have distinct somatosensory demands, so we built the canine DRG atlas using ganglia from spinal segments commonly assessed in clinical exams (L5: femoral, L7: sciatic, S1: sciatic/perineal/urination/defecation). We determined whether specific cell types were differentially represented across spinal segments using only the 10x Genomics data to control for possible technical artefacts. Limited sample size motivated collapsing neuronal subtypes into established classes (Ab-LTMR, Ad-LTMR, A-PEP, C-LTMR, C-PEP, C-NP, and C-Thermo), followed by Wilcoxon rank-sum testing between pseudoreplicates of lumbar (L5 and L7) and sacral DRGs for each class. Ab-LTMRs were significantly enriched in lumbar DRGs compared to sacral DRGs (6.3% higher; adjusted P < 0.0001), whereas A-PEPs were significantly enriched in sacral DRGs (4.2% higher; adjusted P < 0.0001; **Figure 4A**). Within the non-neuronal population, non-myelinating Schwann cells and endothelial cells were enriched in lumbar DRGs, while macrophages were enriched in sacral DRGs (**Figure 4B**). While these observations are based on a limited number of canines (n = 3), the observed neuronal and non-neuronal shifts may reflect known differences in DRG function along the spinal axis, including distinctions between limb mechanosensation and enteric or pelvic organ sensitivity [51; 55].

**Figure 4:**
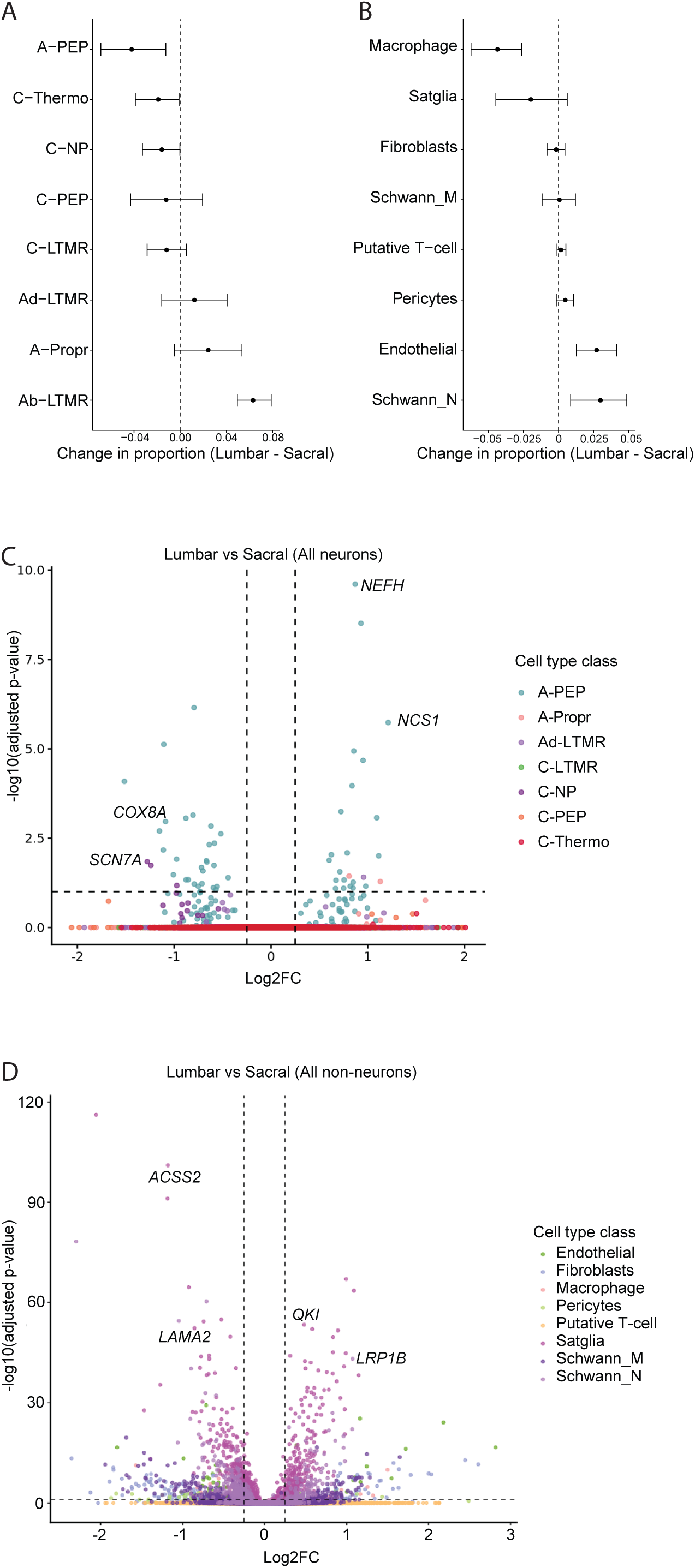
Variation in canine DRG subtype proportions across spinal levels. A) Box and whisker plot displays the difference in proportion of neuronal classes between lumbar and sacral DRG. Data were calculated by resampling the data 10 times (bootstrapping; see Methods) and whiskers represent confidence intervals across resamples. B) Box and whisker plot displays the difference in proportion of non-neuronal subtypes between lumbar and sacral DRG. Data were calculated by resampling the data 10 times (bootstrapping; see Methods) and whiskers represent confidence intervals across resamples. C) Volcano plot displays genes differentially expressed in lumbar DRG compared to sacral DRG (log2FC > 0.5; adjusted p-value < 0.05; dotted line) for each neuronal subtype. Dots are colored neuronal subtype classes. D) Volcano plot displays genes differentially expressed in lumbar DRG compared to sacral DRG (log2FC > 0.5; adjusted p-value < 0.05; dotted line) for each non-neuronal subtype. Dots are colored neuronal subtype classes.

Spinal segment-specific transcriptional differences may also contribute to functional specialization, so we compared subtype-specific gene expression between lumbar and sacral DRGs. We analyzed differential gene expression separately for each neuronal subtype, contrasting lumbar to sacral DRGs. Across all subtype-level tests, we identified 56 differentially expressed genes in neuronal subtypes (|log2FC| > 0.5; adj. p-value < 0.1; **Figure 4C; Table S4**) and 1,089 differentially expressed genes in non-neuronal subtypes (|log2FC| > 0.5; adj. p-value < 0.1; **Figure 4D**). Notably, 50 of the 56 genes in the neuronal subtypes arose from the A-PEP.KIT and A-PEP.NTRK3 contrasts, and this pattern persisted after controlling for differences in cell numbers in the differential expression analyses (**Table S5**). Similar to the subtype proportion differences, the pronounced lumbar–sacral transcriptional shifts in A-PEP neurons may reflect increased sensitivity required for their innervation target, such as reproductive organs.

### Canine DRG cell subtypes are broadly similar to other mammalian subtypes

Comparing transcriptomes between human and canine DRG remains critical for translational insights across species. To compare cell subtypes between canines and humans, we performed two orthogonal analyses on the recent NIH PRECISION Pain human reference atlas [6]: (1) we integrated the canine reference atlas with the human reference atlas to cluster human and canine atlases together and (2) we used a label-transfer approach trained on the human reference atlas to assign human cell types to dog clusters (anchoring) [52]. We evaluated transcriptomic relationships in the integrated human–canine dataset using hierarchical clustering based on Euclidean distances between cluster centroids of the top 30 principal components and label transfer based on prediction scores and the proportion of query cells assigned to a reference cell type.

Across both analytical approaches, we identified consistent transcriptomic relationships between human and canine DRG subtypes (**Figure 5A and 5B**). For example, hierarchical clustering of the integrated human–canine dataset grouped human C-LTMR.CDH9 most closely with dog C-LTMR.CDH9. Similarly, the label transfer approach assigned ∼99% of dog C-LTMR.CDH9 with human C-LTMR.CDH9. However, certain subtypes had a many-to-one relationship between humans and dogs. For example, the human reference atlas has three proprioceptor subtypes, but independently clustering the canine data yielded only two proprioceptor populations. The canine A-Propr nuclei integrated and anchored with one of the three human proprioceptor populations. The lack of granularity in the canine data is likely a consequence of the relative low number of nuclei sequenced compared to humans [6; 7].

**Figure 5:**
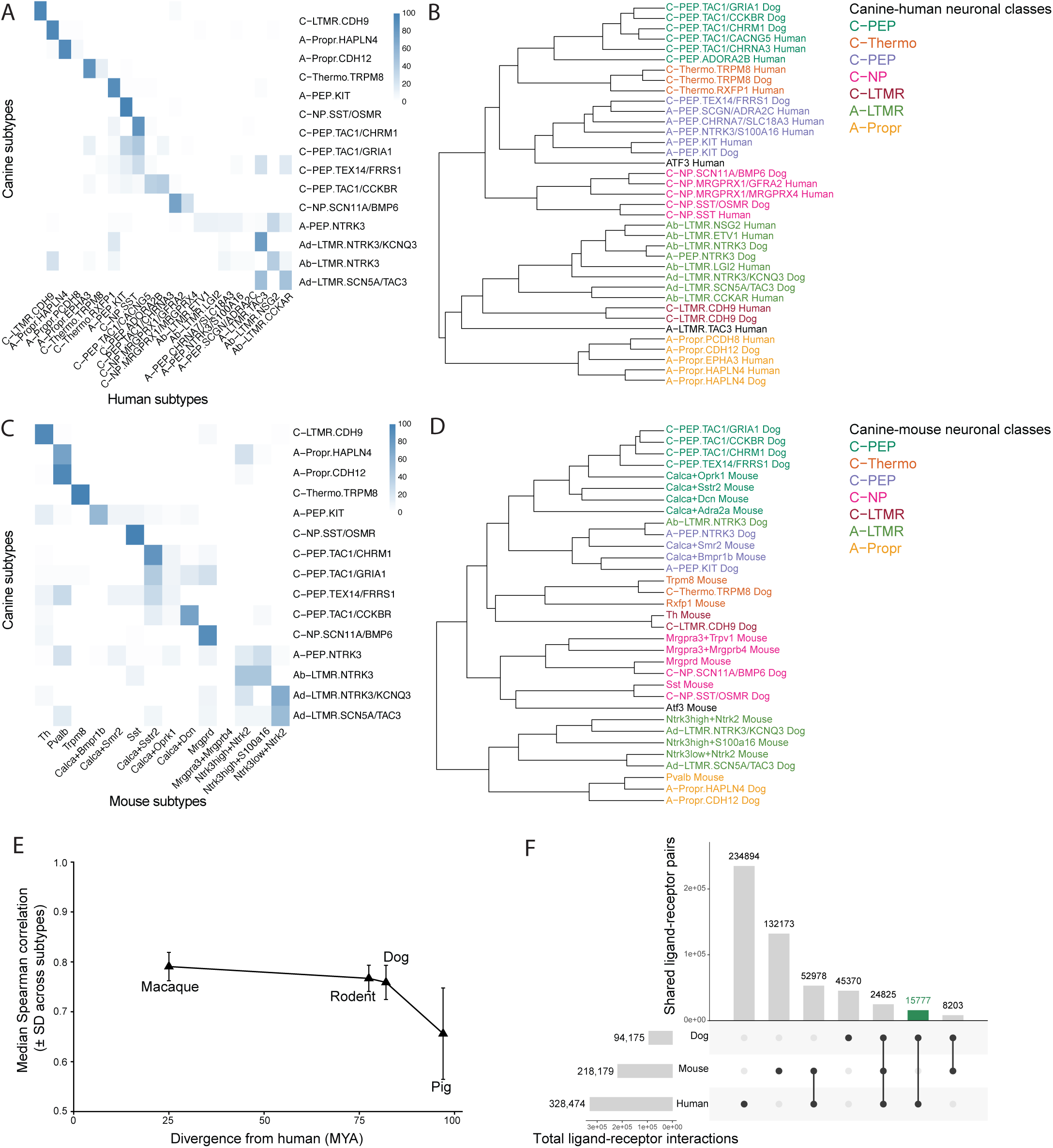
Concordance between Dog, Human, and Mouse Atlases. A) Heatmap displays the percentage of predicted human subtypes for each canine subtype. Percentages were determined after anchoring the canine atlas to the recent human reference atlas using the Seurat lab transfer approach.[6] Only cells/nuclei with an anchoring score > 0.5 are presented. B) Dendrogram displays cell type similarities of canine and human DRG neurons after integration of the canine reference atlas and the recent human reference atlas. [6] This hierarchical relationship was calculated based on the Euclidean distance of the principal components. Branch lengths are scaled to Euclidean distance. C) Heatmap displays the percentage of predicted mouse subtypes for each canine subtype. Percentages were determined after anchoring the canine atlas to a mouse reference atlas, updated from the mouse data in Bhuiyan et al., 2024 (See Methods). Only cells/nuclei with an anchoring score > 0.5 are presented. D) Dendrogram displays cell type similarities of canine and mouse DRG neurons after integration of the canine reference atlas and a mouse reference atlas, updated from the mouse data in Bhuiyan et al., 2024 (See Methods). This hierarchical relationship was calculated based on the Euclidean distance of the principal components. Branch lengths are scaled to Euclidean distance. E) Transcriptomic correlation of DRG cell types to humans decreases over evolutionary distance. For each species, the correlation between the average expression of all genes (log-normalized counts) in each cell type to the average expression of all genes in the corresponding human cell type (y axis; triangle, median correlation; error bars, standard deviation), plotted against the evolutionary distance from the last common ancestor with humans [million years (Ma) ago; x axis]. Macaque species and rodent species (mouse, rat and guinea pig) were grouped together. F) UpSet plot displays the number of overlapping and species-specific predicted ligand–receptor interactions across canine, human, and mouse datasets. The bar plot on the left shows the total number of predicted ligand–receptor interactions in each species. The bar plot on the top shows the number of interactions for each specific overlap category indicated by the connected dots in the matrix below.

The reliance on murine studies for our fundamental understanding of DRG cell type function necessitates a comparative analysis of canine and mouse transcriptomes. We approached our comparison between canine and murine DRG subtypes similarly to our comparison between canine and humans: (1) we integrated the canine reference atlas with the human reference atlas to cluster murine and canine atlases together and (2) we used a label-transfer approach trained on the murine reference atlas to assign human cell types to dog clusters. The murine atlas that we used is an updated version of the harmonized DRG atlas [6; 7]. Again, across both analytical approaches, we identified consistent transcriptomic relationships between mouse DRG subtypes and canine DRG subtypes (**Figure 5C** and **5D**).

While canine subtypes generally corresponded to the same mouse or human subtype, there were two key exceptions: canine A-PEP.NTRK3 and Ab-LTMR.NTRK3. In our human analyses, both subtypes mapped to the LTMR class; however, in mice, both mapped to the A-PEP class. This pattern may signal a biological divergence in canine A -fibers from mouse and human, although technical artifacts cannot be excluded. Importantly, both populations are represented across sequencing platforms and multiple libraries and persist across multiple QC thresholds. Together, these observations suggest that the discrepancy is robust and warrants targeted validation to distinguish biological signal from residual technical effects.

Beyond mouse and human DRG studies, the canine DRG atlas contributes to a growing body of literature surrounding the DRGs from non-traditional model organisms. We also compared cell-type-specific transcriptomic profiles of sensory ganglia from dogs, human, nonhuman primates, mouse, rats, guinea pig, pigs, and axolotls by calculating the correlation between the average normalized counts for each corresponding cell type between each species (**Figure 5E**). Consistent with prior reports, we found that, as evolutionary distance from humans increased, the transcriptomic correlation with the most similar non-human sensory cell type decreased.

Canines offer many benefits in drug development compared to mice, including their human-like metabolism and larger body size [24; 33; 34]. Thus, we next compared predicted ligand–receptor interactions across species, which can help identify conserved drug targets, to evaluate the strengths and weaknesses of dogs as a translational model for pain (**Figure 5F**). To do this, we performed our LR analyses again, but this time on the predicted human cell type for the canine and murine neurons (**Figure 5F; Table S6**). These LR analyses reveal 15,777 predicted ligand–receptor interactions that are found in dog and human but not in mice. Notably, the dog–human specific interactions span pathways with established therapeutic relevance, including NGF signaling through NTRK1 (49 predicted interactions) and NGFR (78 predicted interactions) as well as signaling involving GFRA1 (34 predicted interactions) and RET (22 predicted interactions) [54]. Together, these results suggest that dogs capture a substantial set of potentially druggable intercellular signaling programs shared with humans that are underrepresented in mice, supporting their value as a complementary translational model for pain.

## Discussion

Pain is a common clinical complaint in dogs [39; 40]. Causes of pain include but are not limited to musculoskeletal, neurologic, reproductive, and oncologic pain [42]. However, there are insufficient treatments available for pain management in dogs, and pain is underrecognized, undertreated, and is frequently reported as a cause of euthanasia [38]. A key gap in the field is the limited understanding of the molecular features of canine pain-sensing sensory cell types. In this study, we generated a canine DRG cell atlas to define those molecular features and to provide a foundation for the development of new pain therapies for dogs with potential relevance for translation to human pain biology and treatment.

To address the limited understanding of the molecular features that define canine pain-sensing sensory cell types, we identified fifteen neuronal and eight non-neuronal cell types in canine DRG and assigned them to established classes based on gene expression. To place these cell types in a broader comparative framework, we aligned our nomenclature with the human DRG atlas, consistent with terminology recently adopted by the NIH PRECISION Human Pain Network [6]. Cross-species comparison with published human and mouse neuronal atlases revealed that the canine atlas contains counterparts to each major neuronal subtype identified in those species. Differences likely reflected the smaller number of canine nuclei relative to the human and mouse datasets, which limited our ability to resolve finer neuronal subpopulations in dogs. Together, these findings establish the canine DRG atlas as a resource for understanding the molecular organization of sensory cell types in dogs and for guiding the development of therapies with potential relevance to both canine and human pain biology. To support future studies, we made both the neuronal and non-neuronal atlases publicly available through the Painseq web resource: http://painseq.shinyapps.io/canine_drgs/.

Dogs offer several advantages as a translational model for pain research that extend beyond molecular similarity. Unlike induced rodent models, dogs develop naturally occurring pain conditions, including orthopedic and neuropathic disorders, in the context of an intact immune system, complex environment, and heterogeneous genetic background [31; 41; 48]. This may better reflect the variability seen in human pain populations. Dogs also share environmental exposures and social contexts with humans, and their larger body size enables the use of clinical imaging modalities, devices, and dosing strategies that are more difficult to model in mice [30; 53]. Together, these features make dogs a valuable complementary model for evaluating pain mechanisms and therapeutic responses in settings that may more closely approximate human disease.

We performed LR, or cell-cell communication, analyses using LIANA to guide prioritization of therapeutic targets [12]. Once we correlated cell types across species, we identified 94,175 LR interactions in dog, 218,179 in mouse, and 328,474 in human DRG datasets. The smaller number detected in dog DRG likely reflects both more limited genome annotation in dog relative to human and mouse DRG and the fewer canine cells/nuclei [23]. Despite this lower total, we identified 15,777 LR interactions that were shared between dog and human but not mouse DRG. This suggests that dogs may capture aspects of DRG cell-cell communication that are more similar to humans than those represented in mice. These shared interactions may be especially informative for evaluating the efficacy and potential side effects of new treatments. For example, we identified predicted interactions involving *NTRK1* and *NGFR* in dogs and humans that were not detected in mice. Such shared LR interactions may also help identify compounds or treatment strategies for reverse translation, in which medications with known mechanisms or efficacy in humans are evaluated for use in dogs with chronic pain [16].

One clinically relevant pathway example is NGF represented by signaling. Bedinvetmab (Librela), an anti-nerve growth factor antibody, has been rapidly adopted in veterinary medicine for the treatment of chronic pain [9; 36]. NGF-receptor interactions normally support neuronal survival and function, but blocking this pathway may reduce nociceptive signaling in the DRG [35]. We identified the neuronal cell types that express NGF receptors, including TrkA and p75, as these populations may be those most directly affected by Bedinvetmab treatment [46]. Although Tanezumab, the humanized anti-NGF antibody, was ultimately not approved for human use, canine and feline anti-NGF therapies are currently in clinical use [14]. These interspecies differences highlight the need for further investigation into the functional and safety consequences of modulating NGF signaling across species.

In addition to cross-species comparisons, our dataset enabled us to examine variation across spinal levels within dogs. We generated libraries from individual DRG and identified 98 to 819 neurons per ganglion. Although neuronal yield varied by DRG segment, it also improved over the course of the study, as we optimized the microdissection of the ganglia. We selected L5, L7, and S1 because these levels contribute to distinct named nerves in the dog [3; 8]. In most dogs, L7 and S1 contribute to the sciatic nerve, whereas L5 contributes variably to the tibial and common peroneal branches of the sciatic nerve as well as the femoral nerve. The S1 level additionally contributes to the dorsal nerve of the penis in males and the superficial perineal nerves [50; 51]. These levels have clinical significance in dogs, as they may be affected intervertebral disc disease, lumbosacral stenosis, or other disorders of the spinal cord and nerve roots and damage to these sensory levels can result in paraparesis or urinary and fecal incontinence. While multiple human and mouse DRG datasets are available for lumbar levels, comparable sacral DRG datasets are currently unavailable. As a result, future studies will be needed to determine how sacral canine DRG compares with sacral DRG in humans and mice.

Comparison of lumbar and sacral DRGs provided an opportunity to explore regional differences in the relative representation of afferents. Within our limited sample of three dogs, we observed proportional shifts across several neuronal classes. Specifically, we noted an enrichment of Ab-LTMRs in the lumbar DRG, alongside an enrichment of A-PEP neurons in the S1 DRG that was driven primarily by the A-PEP.KIT subpopulation. Ab-LTMRs are commonly associated with the innervation of haired skin [45]. Furthermore, A-PEP.KIT neurons correspond to A-HTMRs, a class implicated in colonic as well as cutaneous innervation [19; 55]. We therefore hypothesize that the relative abundance of Ab-LTMRs in the lumbar DRG may reflect a greater representation of haired skin afferents. Conversely, the increased proportion of A-PEP neurons in the sacral DRG could indicate specialized innervation of internal organs or pelvic functions related to urination and defecation. Together, these preliminary findings suggest potential functional specialization along the spinal axis, though further anatomical tracing is required to definitively confirm these innervation targets [51].

Several technical features of our study should be considered when interpreting these findings. We performed single-nucleus sequencing for the 10x datasets because the timing of donor DRG collection was unpredictable, requiring tissues to be frozen at −80°C between collection and nuclei isolation. Nuclear RNA-seq can influence gene expression measurements, including the proportion of intronic reads and the total number of genes detected [13]. However, profiling nuclei rather than whole cells also offers important advantages, including reduced do against large cells during library preparation and the ability to batch-process samples collected over time [25]. The combination of Flash-seq and 10x data was, therefore, a strength of this study. The 10x datasets provided broad cellular coverage, enabling detection of rare cell types such as C-THERMO.RXFP1 and C-LTMR.CDH9, whereas Flash-seq provided deeper sequencing coverage that increased confidence in neuronal identification [13].

In the 10x sequencing datasets, neurons comprised a relatively high proportion of nuclei, ranging from 10% to 40% depending on the individual library. This variability may reflect differences in tissue harvesting, individual dogs, spinal level, or other technical factors. We also made a concerted effort to trim as much connective tissue as possible during dissection, which may have contributed to neuronal enrichment. Previous studies of DRG have likewise reported substantial variability in neuron-to-non-neuron ratios, with estimates ranging from 5% to 60% depending on species and methods of tissue collection and processing [25; 46].

## Conclusion

This atlas provides a framework to understand the canine DRG, including spinal level and cross-species comparisons. Pain management in dogs and humans continues to be a challenging clinical problem. In dogs, diagnosis of pain is challenging, with variable methods based on clinical exam, owner description of the behavior, imaging, or treatment trials. Additionally, for humans, there is not a single treatment for pain that is universally effective, and in many cases, treatments have side effects that are not benign. The lack of current effective treatments indicates that the dog and human may both benefit from continued development of pain treatments. We establish several cell types in the canine DRG, report on the similarities and differences between dogs, humans, mice, and other organisms, and provide a resource of information to be used for future drug development for canine and human pain research.

## Data availability

Processed data are available at https://painseq.shinyapps.io/canine_drgs/. Raw and processed data will be available within the Gene Expression Omnibus (GEO:GSEXXXXXX) repository (www.ncbi.nlm.nih.gov/geo).

## Acknowledgements

We would like to thank the donors and the donors’ families, without whom much of this study would not be possible. We would like to thank Paula Ledesma Fernandez, Greg Weir, and Andrew Bell from the University of Glasgow for generously sharing their dataset and their collegial assistance during this process. We would also like to thank members of the Renthal and Parker labs as well as A. Salman, M. Xu, and A. Shuster for helpful feedback throughout the study. Thank you to Dr. Hehuang David Xie for the generous use of his space and equipment. This research was funded through the Internal Research Competition at the VA-MD College of Veterinary Medicine to RLP, AB, and TC. W.R. was supported by the National Institute of Neurological Disorders and Stroke [U19NS130617, R01NS119476]. W.R. receives additional support from the Burroughs Wellcome Fund, Rita Allen Foundation, National Institute of Drug Abuse (DP1DA054343), National Eye Institute (U01EY034709), BWH Women’s Brain Initiative, BWH Neurotechnology Studio, and MGB Gene and Cell Therapy Institute.

## Conflict of Interest

The authors declare that the research was conducted in the absence of any commercial or financial relationships that could be construed as a potential conflict of interest.

## Notes

https://painseq.shinyapps.io/canine_drgs/

### Competing Interest Statement

The authors have declared no competing interest.

### Summary of Updates

Update the reference and add the DOI for the bioRxiv version of the FLASH-seq data by Fernandez et al.

