## Supplementary Figures, Table S1, Legends for "Translational insights into canine dorsal root ganglia cell types using cross-species comparisons"

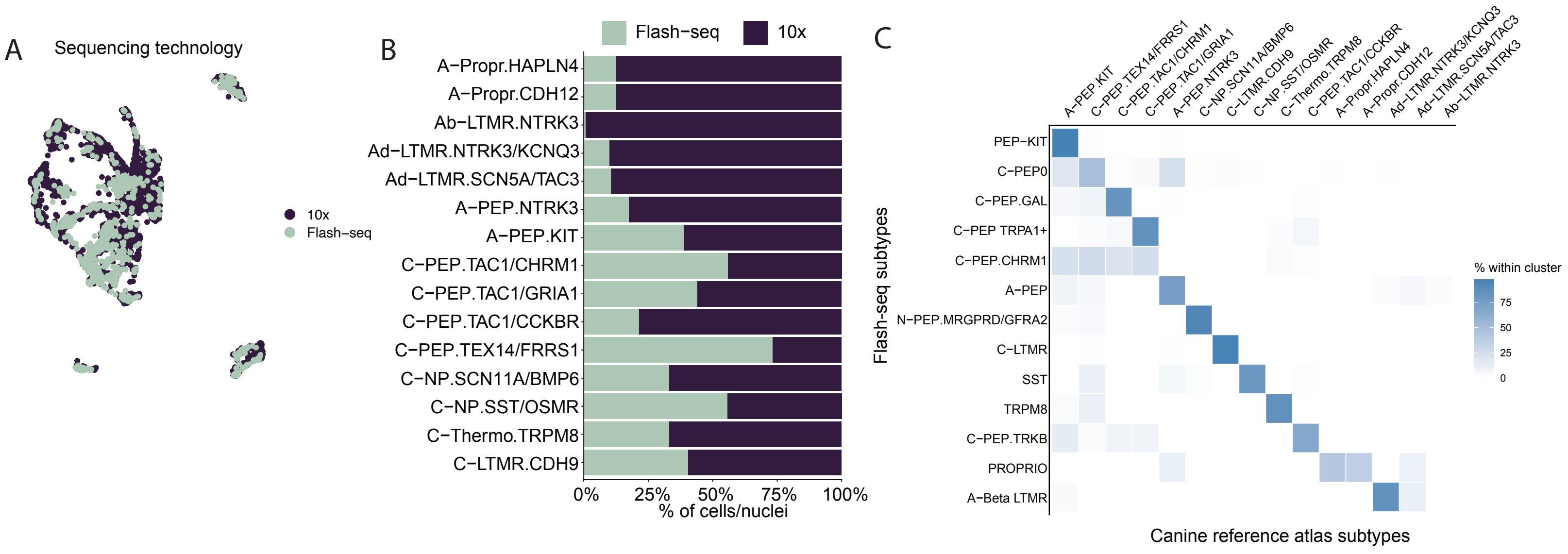

A
Sequencing technology
10x
Flash−seq
B
Flash−seq
10x
A−Propr.HAPLN4
A−Propr.CDH12
Ab−LTMR.NTRK3
Ad−LTMR.NTRK3/KCNQ3
Ad−LTMR.SCN5A/TAC3
A−PEP.NTRK3
A−PEP.KIT
C−PEP.TAC1/CHRM1
C−PEP.TAC1/GRIA1
C−PEP.TAC1/CCKBR
C−PEP.TEX14/FRRS1
C−NP.SCN11A/BMP6
C−NP.SST/OSMR
C−Thermo.TRPM8
C−LTMR.CDH9
0%
25%
50%
75%
100%
% of cells/nuclei
C
A−PEP.KIT
C−PEP.TEX14/FRRS1
C−PEP.TAC1/CHRM1
C−PEP.TAC1/GRIA1
A−PEP.NTRK3
C−NP.SCN11A/BMP6
C−LTMR.CDH9
C−NP.SST/OSMR
C−Thermo.TRPM8
C−PEP.TAC1/CCKBR
A−Propr.HAPLN4
A−Propr.CDH12
Ad−LTMR.NTRK3/KCNQ3
Ad−LTMR.SCN5A/TAC3
Ab−LTMR.NTRK3
Flash-seq subtypes
PEP−KIT
C−PEP0
C−PEP.GAL
C−PEP TRPA1+
C−PEP.CHRM1
A−PEP
N−PEP.MRGPRD/GFRA2
C−LTMR
SST
TRPM8
C−PEP.TRKB
PROPRIO
A−Beta LTMR
% within cluster
75
50
25
0
Canine reference atlas subtypes

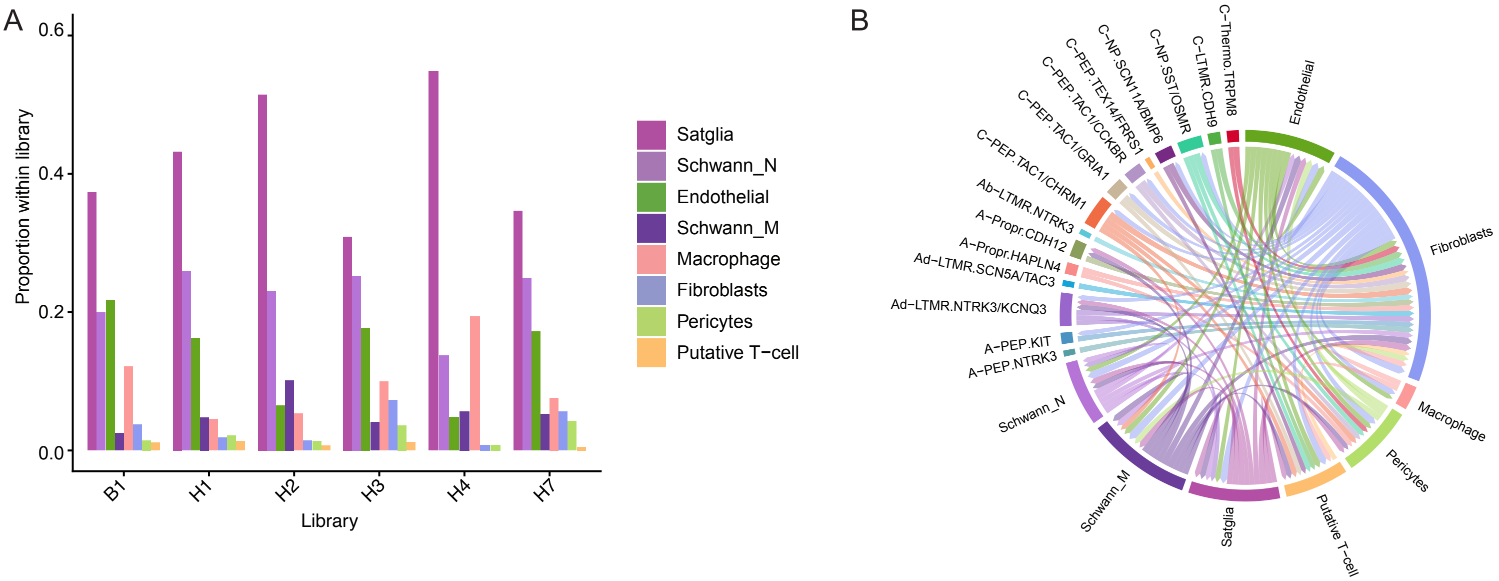
A
B
0.6
0.4
0.2
0.0
Proportion within library
B1
H1
H2
H3
H4
H7
Library
Satglia
Schwann_N
Endothelial
Schwann_M
Macrophage
Fibroblasts
Pericytes
Putative T−cell
C−Thermo.TRPM8
C−LTMR.CDH9
C−NP.SST/OSMR
C−NP.SCN11A/BMP6
C−PEP.TEX14/FRRS1
C−PEP.TAC1/CCKBR
C−PEP.TAC1/GRIA1
C−PEP.TAC1/CHRM1
Ab−LTMR.NTRK3
A−Propr.CDH12
A−Propr.HAPLN4
Ad−LTMR.SCN5A/TAC3
Ad−LTMR.NTRK3/KCNQ3
A−PEP.KIT
A−PEP.NTRK3
Schwann_N
Schwann_M
Satglia
Putative T−cell
Pericytes
Macrophage
Fibroblasts
Endothelial

**Table S1**

| Sample | Library | Dog | Spinal Level | Number of Neurons | Number of Non-neurons |
|---|---|---|---|---|---|
| 1 | A1 | B | L5 | Excluded | Excluded |
| 2 | B1 | C | L5 | 277 | 882 |
| 3 | H1 | B | L7 | 98 | 375 |
| 4 | H2 | C | L7 | 226 | 5349 |
| 5 | H3 | A | S1 | 581 | 1934 |
| 6 | H4 | B | S1 | 112 | 124 |
| 7 | H7 | A | L7 | 819 | 3070 |

### Supplementary Figure and Table Legends

*Figure S1: The neuronal canine atlas contains 10x Genomics snRNA-seq and Flash-seq data*

A) UMAP projection displays canine neuronal DRG atlas colored by sequencing technology (3,096 cell/nuclei)

B) Bar plots display the composition of sequencing technologies for each neuronal subtype. Bars are colored by sequencing technology.

C) The heatmap displays the percentage of subtype annotations in the canine DRG neuronal atlas that correspond to those independently identified in the Flash-seq datasets.

*Figure S2: Reproducibility of the canine non-neuronal reference atlas*

A) Bar plots display the proportion of non-neuronal subtypes per sequencing library.

Table S1: Source of DRG for 10x sequencing experiments.

B) Chord diagram displays ligand–receptor predicted interactions between neuronal and non-neuronal subtypes after downsampling each subtype to 100 cells/nuclei. Only subtypes with an aggregated rank < 0.01 are displayed.

**Tables:**

Table S1: Description of the libraries generated in the 10x dataset. Seven libraries were generated but data from 6 was included in the data set. Dog A was a 2 year-old Female spayed Rottweiler that was euthanized for osteosarcoma in the thoracic limb, dog B was a 3 year-old Male Intact Pitbull that was euthanized for a behavioral condition, and dog C was an approximately 5 year-old Female Intact Pitbull mix that was euthanized for unknown causes. The library name, dog, spinal level, and nuclei yield are reported.

Table S2: Neuronal marker genes generated using Seurat's FindAllMarkers function. Test of differential expression was performed between each neuronal subtype and all other neurons.

Table S3: Non-neuronal marker genes generated using Seurat's FindAllMarkers function: Test of differential expression was performed between each non-neuronal subtype and all other non-neurons.

Table S4: Subtype- and spinal region-specific marker genes generated using Seurat's FindAllMarkers function. Test of differential gene expression was performed between each subtype at each level and the same subtype at all other levels.

Table S5: Ligand receptor interactions between canine neurons and non-neurons

Table S6: Shared and species-specific ligand receptor interactions between neurons and non-neurons in canines, mice and humans

Table S1: Source of DRG for 10x sequencing experiments.

Table S1: Source of DRG for 10x sequencing experiments.
